# Targeted finetuning enables co-folding models to learn ligand-induced protein conformational states

**DOI:** 10.64898/2026.09.21.752570

**Authors:** Rohan Gorantla, Christian Schleberger, Fabian Sesterhenn

## Abstract

Advances in protein structure prediction have enabled all-atom protein-ligand co-folding models that predict bound conformations directly from sequence and small-molecule structure. However, these models often fail to generalize to novel binding sites or alternative protein conformational states, limiting their utility for chemical biology and drug discovery. Here we show this limitation reflects training data bias rather than architectural constraints and can be overcome through targeted finetuning. Using ten previously unseen X-ray structures of Werner (WRN) helicase from a drug discovery program, we finetune Boltz-1 to learn both an allosteric binding site and a large conformational change locking the enzyme in an inactive state, while preserving accuracy on the ATP-bound state. The finetuned model generalizes to different chemical series and transfers the conformational logic across RecQ-family helicases in a binding-site sequence-dependent manner. This approach provides a blueprint for adapting foundation models as new structural and mechanistic data emerge, enabling co-folding networks to capture ligand-induced conformational switches and binding poses absent from their training data but central to biological regulation and therapeutic intervention.

## Introduction

Deep learning-based protein structure prediction has reshaped structural biology, with models such as AlphaFold achieving high accuracy across a wide range of protein targets^1,2^. These advances have accelerated the integration of predicted structures into biological research and motivated the development of protein–ligand co-folding methods that aim to model ligand-bound states directly from protein sequence and chemical structure.

The release of AlphaFold 3 extended structure prediction to the joint modelling of proteins, nucleic acids, and small molecules within a unified framework^1^. Open-source co-folding models, including Boltz-1^3^, Chai-1^4^, and Protenix ^5^, have further broadened access to these capabilities by providing publicly available model weights and inference pipelines. Together, these methods have raised expectations that such models could support ligand design, binding-mode and affinity prediction, and conformational hypothesis generation in chemical biology and drug discovery ^6^.

Despite this progress, recent benchmarks indicate that current models remain limited in their ability to predict ligand-induced conformational changes. Induced fit, allostery, cryptic-pocket formation, and alternative domain arrangements are sparsely represented in public structural databases relative to canonical apo and holo states. As a result, these models may preferentially reproduce familiar conformations and binding-site geometries, limiting their generalization to underrepresented modes of molecular recognition ^7,8^.

Werner syndrome helicase (WRN) is a challenging example that combines several of these limitations. WRN is a multidomain RecQ-family enzyme and has emerged as a promising synthetic-lethal target in microsatellite instability-high cancers^9^. The first structure of the WRN helicase core, bound to ADP and representing an active conformation, was released in 2020^10^. More than 25 additional structures were released between 2024 and 2026 and revealed an inactive conformation involving a large domain rotation induced by small-molecule inhibitors binding to an allosteric pocket. One of most notable inhibitors is HRO761 (PDB ID 8PFO), which binds at the interface of the D1 and D2 helicase domains to lock WRN in an inactive conformation, and has entered human clinical trials^11^. Given the Boltz-1 training cutoff was 2021, the model had limited exposure to both this ligand-binding pocket and the associated conformational transition. Here, we evaluate whether targeted finetuning on unseen inhibitor-bound structures can improve a foundational model’s ability to capture allosteric ligand binding and large-scale conformational change, providing a practical route for extending such models to settings where conformational dynamics and local interactions determine binding and function.

## Results

### Co-folding models fail to capture ligand-induced conformational state of WRN

We first evaluated the performance of publicly available protein–ligand co-folding models (Boltz-1, Boltz-2^12^, Protenix, and Chai) on WRN helicase in complex with either ATPγS or the inhibitor HRO761, which locks WRN in an inactive conformation (Fig 1A). In zero-shot mode, all models accurately reproduced the overall fold and domain arrangement for the apo and ATP-bound states. However, none recovered the binding mode or the large-scale conformational change induced by HRO761. Predicted complexes remained biased toward the active, ATP-like arrangement regardless of the ligand input (Fig. 1B). The closed D1-D2 interface and ∼180° domain rotation characteristic of the HRO761-bound crystal structure failed to form in any prediction, and ligand poses were frequently mispositioned or sterically clashed with the incorrectly open conformation. These observations confirm that, under zero-shot conditions, off-the-shelf co-folding models default to conformations prevalent in their training data (e.g., nucleotide-bound states) and lack the ability to infer novel ligand-dependent allosteric transitions.

**Fig. 1:**
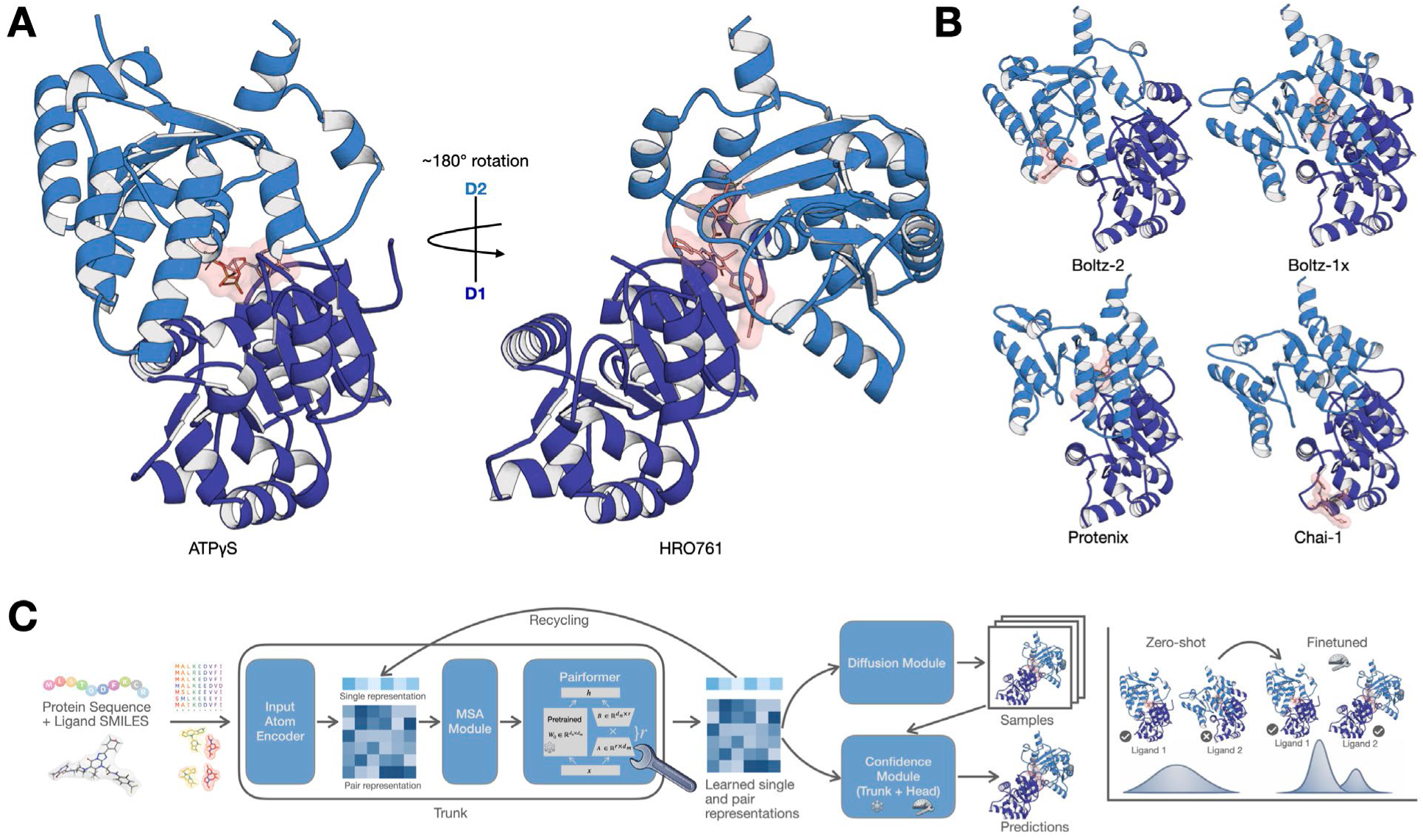
Baseline co-folding models fail to predict the inhibitor-induced WRN conformation under zero-shot conditions. **A.** Crystal structures of WRN helicase bound to ATPγS (active state) and inhibitor HRO761 (inactive state, PDB 8PFO), highlighting the ∼180° D2-domain rotation and closed D1–D2 interface induced by HRO761. **B.** Zero-shot predictions for HRO761-bound WRN by pre-trained co-folding models (Boltz-1, Boltz-2, Protenix and Chai) all adopt an ATP-like open conformation, failing to form the closed interdomain pocket or correctly position the ligand. **C.** Schematic of the co-folding model architecture and finetuning approach: a diffusion-based protein–ligand co-folding model (Boltz-1) is augmented with low-rank adapters (LoRA, rank r = 8) in pairwise interaction layers and finetuned on ligand-bound WRN structures, shifting its prediction distribution from the active-state bias (zero-shot) towards the HRO761-bound inactive conformation (finetuned).

### Parameter-efficient finetuning recovers the HRO761-bound conformation and key interactions

We next finetuned the Boltz-1 model (with training cut off September 2021) using a small set of non-public WRN structures. Throughout the WRN drug discovery program that led to the clinical candidate HRO761, we have determined ∼95 structures over 6 years to guide the lead optimization and obtain detailed insights into the allosteric pocket and the structural determinants of WRN inhibition (Fig. S1). While these structures are not published, their overall protein conformation is very similar to the published HRO761 structure, but contain chemically diverse ligands. We constructed a training set by performing a time-based split of our data, using the first ten structures for training and HRO761 as a test case. Finetuning was limited to ∼1% of parameters by inserting Low-Rank Adaptation (LoRA)^13^ adapters (rank *r* = 8) into Boltz-1’s pairwise interaction layers (Pairformer blocks), while freezing all other weights. This targeted adaptation enabled accurate predictions of the inhibitor-bound WRN state, as compared to the ground truth structure 8PFO (Fig. 2A). The finetuned model Boltz-FT now captured the full interdomain rotation and correctly formed the cryptic pocket at the D1–D2 interface. The predicted HRO761 binding mode was in close agreement with the crystallographic pose, and specific interactions seen experimentally were recovered, as seen in the hinge loop residues (around Thr728 and adjacent Gly-Phe-Asp) adopted the closed “bend” conformation, an arginine-rich segment on the D2 domain engaged HRO761 within the pocket, and the Walker motif/ATP-binding loop was displaced by ∼10 Å relative to its ATP-bound position, consistent with the domain rotation (Fig. 2A). Crucially, finetuning did not compromise performance on the active state. Boltz-FT maintained high accuracy on ATP-bound WRN, indicating that focusing updates on pairformer layers prevented catastrophic forgetting of the canonical state. To further test the prediction accuracy on a larger dataset with diverse ligands, we tested Boltz-FT on the remaining 75 unseen WRN-ligand complexes (Fig. 2B, 2C). Quantitatively, Boltz-FT outperformed the zero-shot model on held-out inhibitor-bound complexes across all metrics. It achieved low *biSyRMSD* values^14^ (often <2 Å, indicating near-native domain positioning and ligand pose) and high *LDDT-PL* scores (∼0.85–0.9, reflecting accurate binding-pocket geometry). The *LDDT-PLI* metric, which measures recovery of specific protein–ligand contacts, also improved, confirming that the finetuned model captures the correct pattern of interactions rather than simply forcing the ligand into place.

**Fig. 2:**
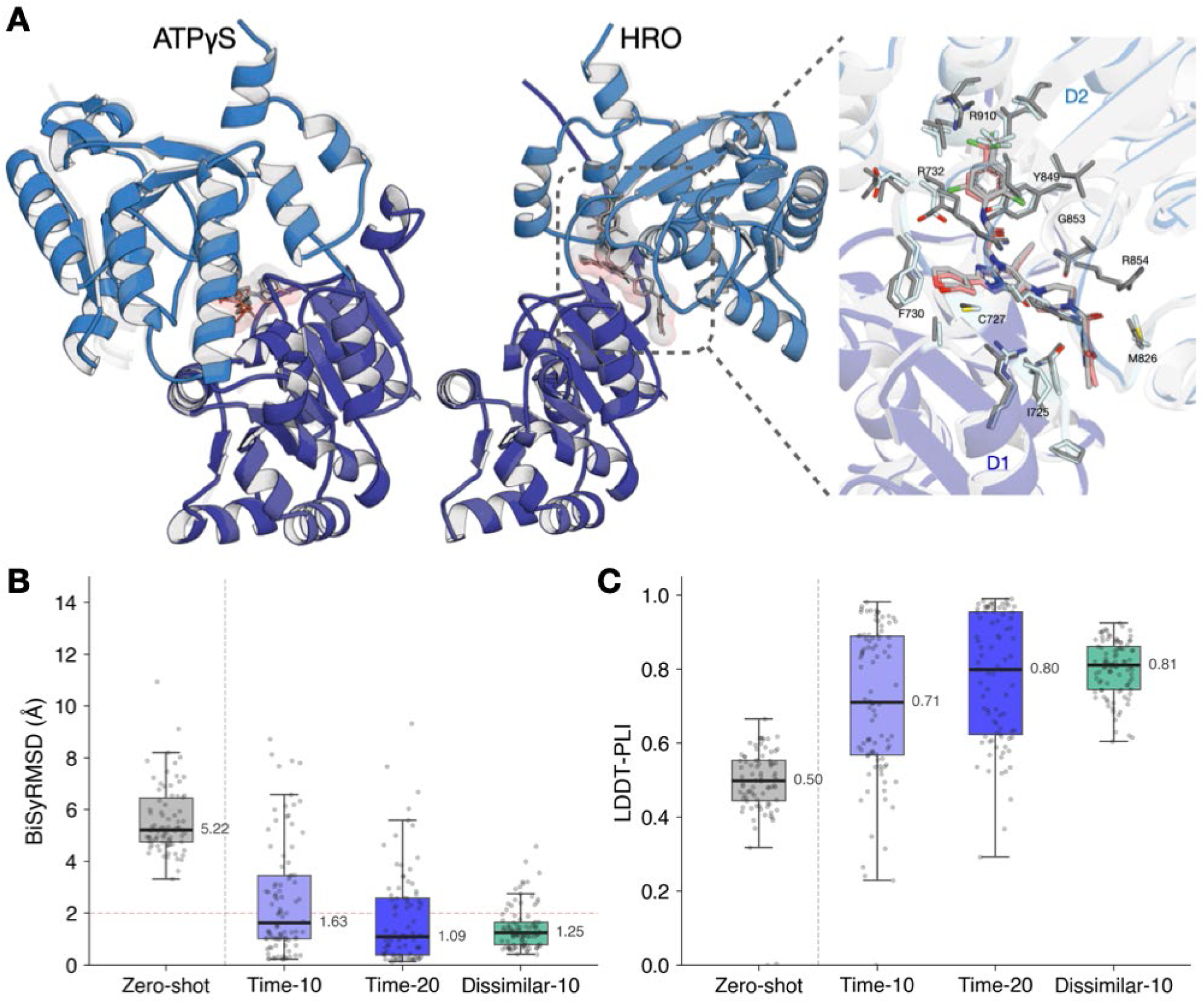
Parameter-efficient finetuning recovers the inhibitor-bound WRN conformation and specific protein-ligand interactions. **A.** Finetuned Boltz-1 (Boltz-FT) accurately predicts the HRO761-bound WRN structure, reproducing the closed D1-D2 domain arrangement and the inhibitor’s binding mode in close agreement with the crystallographic pose (PDB 8PFO). Key interactions in the HRO761-bound state are recovered (e.g., hinge loop around Thr728 adopts the crystallographic closed conformation and an arginine-rich segment in D2 engages HRO761, while the ATP-binding loop is displaced consistent with domain rotation). **B.** Quantitative performance metrics showing that finetuning dramatically improves prediction accuracy on the HRO761-bound complex relative to the zero-shot model. Finetuned models achieve low *biSyRMSD* values (domain-aware symmetric RMSD capturing relative domain orientation and ligand pose accuracy) of ∼1–2 Å versus ∼5 Å for the base model, indicating near-native domain positioning and ligand placement. Time-10 corresponds to 10 training structures from a time-based split, Time-20 to 20 training structures, and Dissimilar-10 to a training set composed of the most dissimilar structures to HRO761 (see Fig S1). **C.** Finetuning also yields higher *LDDT-PLI* scores (quantifying recovery of protein– ligand interactions) of 0.71-0.81 versus 0.50 for the base model, confirming that the finetuned model correctly reproduces the binding pocket geometry and native protein–ligand contacts.

To further investigate the effects of training set choices, we generated a chemical dissimilarity split using ten ligands with lowest Tanimoto similarity (ECFP8 fingerprints) to HRO761 (similarity between 0.1 and 0.2, referred to as Dissimilar-10). These ligands also showed 2-3 orders of magnitude weaker potency in an ATP displacement assay (IC50 between 1.4 µM and 32µM, compared to 8nM for HRO761, Fig. S1). This strategy yielded similar improvements in HRO761 predictions, indicating that mechanistic diversity in the training set and not necessarily the most potent or similar ligand suffices to teach the conformational switch. However, including one ATP-bound WRN structure in the finetuning set was essential, as without any ATP-bound examples, the model overfits to the inactive state and lost the ability to produce the active conformation (i.e., it predicted the closed state even for ATP ligands). With a balanced training set (mixed active and inactive conformations), the finetuned model maintained dual-state competence.

### Boltz-FT predicts conformational changes on independent unseen fragment data

As a further test of generalisation, we evaluated Boltz-FT on WRN–ligand complexes reported by an independent group in 2026^15^.These structures of bound fragment hits and optimized lead compounds capture an inactive WRN conformation (referred as Form D) similar to our HRO761-bound state but were absent from public data during model training. The finetuned model accurately reproduced the closed interdomain arrangement for all Form D test cases (e.g., PDB 9MJU, 9MJV, 9MJY, and 9MJZ), forming the allosteric pocket at the D1-D2 interface and placing each fragment or inhibitor within it. Predicted complexes exhibited minimal domain misalignment relative to the crystal structures (Fig. 3). By contrast, the base model remained biased toward the ATP-bound form, failing to form the closed pocket and mispositioning the ligands in all cases.

**Fig. 3:**
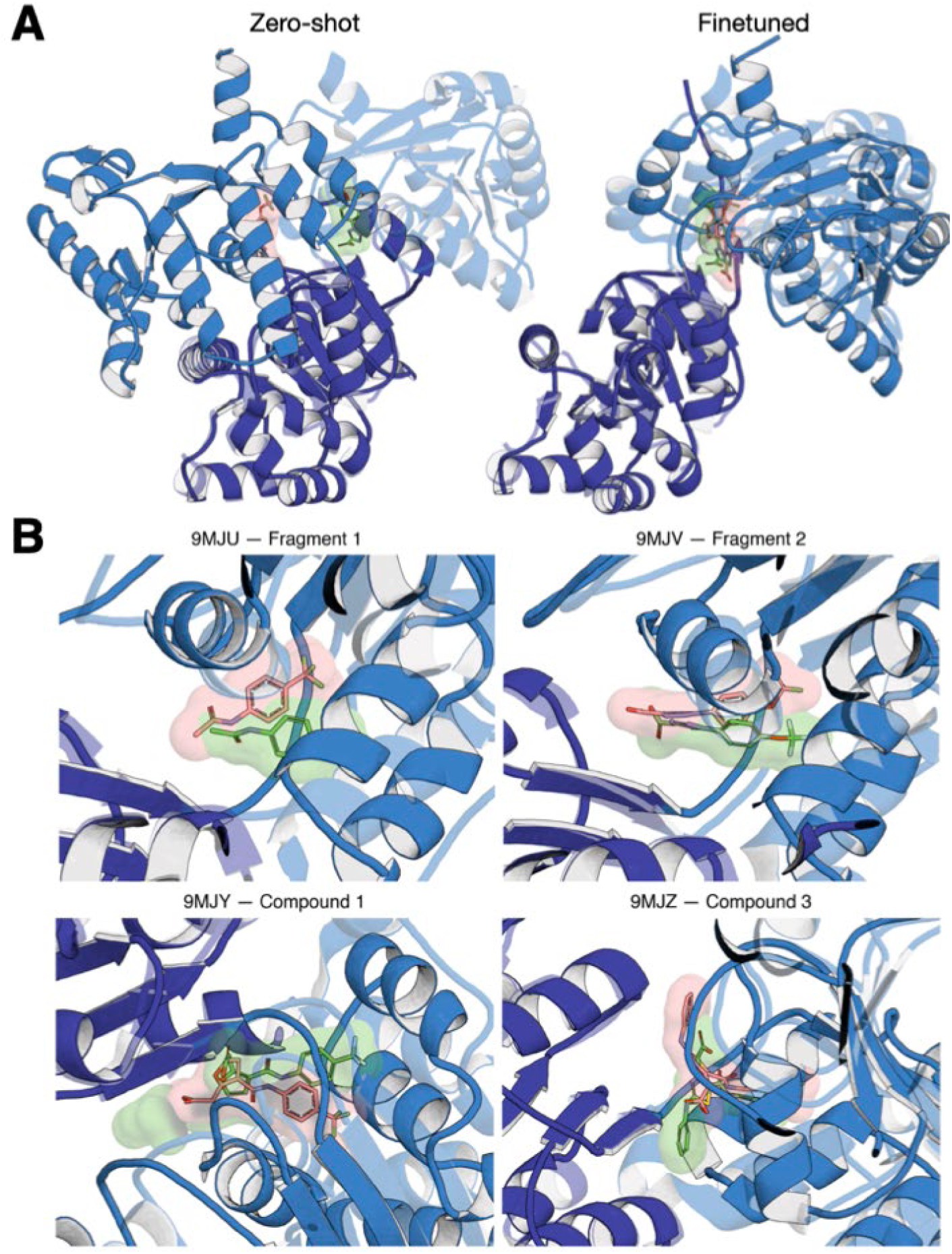
Boltz-FT generalises to external WRN fragment and compound structures and recovers the inactive Form D conformation. **A** Overlay of the crystallographic ground truth structure (green) with zero-shot Boltz-1 and finetuned Boltz-FT predictions for an external WRN-ligand complex from an independent 2026 dataset not used for training. Zero-shot Boltz-1 remains in an open, ATP-like conformation and fails to form the closed D1–D2 allosteric pocket, whereas Boltz-FT recovers the closed interdomain arrangement and places the ligand within the pocket. **B** Close-up views of the D1–D2 allosteric pocket for four external Form D test cases - PDB 9MJU and 9MJV (fragment hits), and PDB 9MJY and 9MJZ (optimized compounds) showing Boltz-FT ligand poses overlaid with the crystallographic ligand positions and pocket geometry.

### LoRA adapter rank controls a tight regularization–capacity tradeoff

We next investigated how the LoRA adapter rank influences model performance, reflecting the trade-off between model capacity and regularization. Increasing the rank from 8 (∼6 million trainable parameters, ∼1% of the full 592M-parameter model) to 16 or 32 (∼2–4% of parameters) did not yield additional performance improvements (Fig. S2). In contrast, a rank of 8 consistently provided strong performance, indicating that this level of capacity is sufficient to capture the relevant conformational changes while maintaining effective regularization. Notably, even under constrained settings, training on chemically distant, weak compounds and limiting the number of trainable parameters, the model successfully generalized to mature, potent compounds such as HRO761. Overall, these ablations demonstrate that few-shot adaptation is robust and does not require extensive parameter updates or highly optimized training data to recover the correct conformational transition.

### Conformational switching logic transfers across the RecQ helicase family showing that local ligand interactions are learned

Finally, we investigated whether the finetuned model’s learned allosteric switch in WRN extends to related helicases (Fig. 4A). Boltz-FT produced ligand-specific predictions for two close homologs, BLM and RecQ1, modeling each in both an active (ATPγS-bound) and an HRO761-bound conformation. The HRO761-bound predictions for BLM and RecQ1 showed the same behaviour as WRN – domain closure and pocket formation – suggesting that the model captured generalizable features of the allosteric mechanism beyond WRN and can learn local interactions between protein and ligand. However, for RecQ5, the finetuned model failed to recapitulate the HRO761-bound conformation; its prediction remained in a state between ATP-like and HRO-like conformation. Sequence analysis revealed that RecQ5 lacks several key binding-site residues present in WRN/BLM/RecQ1 and led us to perform reciprocal mutational tests *in-silico*: introducing six WRN-derived residues into RecQ5’s pocket (RecQ5_C) enabled Boltz-FT to predict the closed, HRO761-bound-like state for RecQ5, whereas introducing the corresponding RecQ5 residues into BLM and RecQ1 (BLM_C, RecQ1_C) prevented the conformational switch (Fig. 4B, Fig. S3). These results indicate that the model’s learned ligand-dependent conformational logic is explicitly sequence-dependent – it triggers the allosteric transition only when the requisite key residue features and interactions are present.

**Fig. 4:**
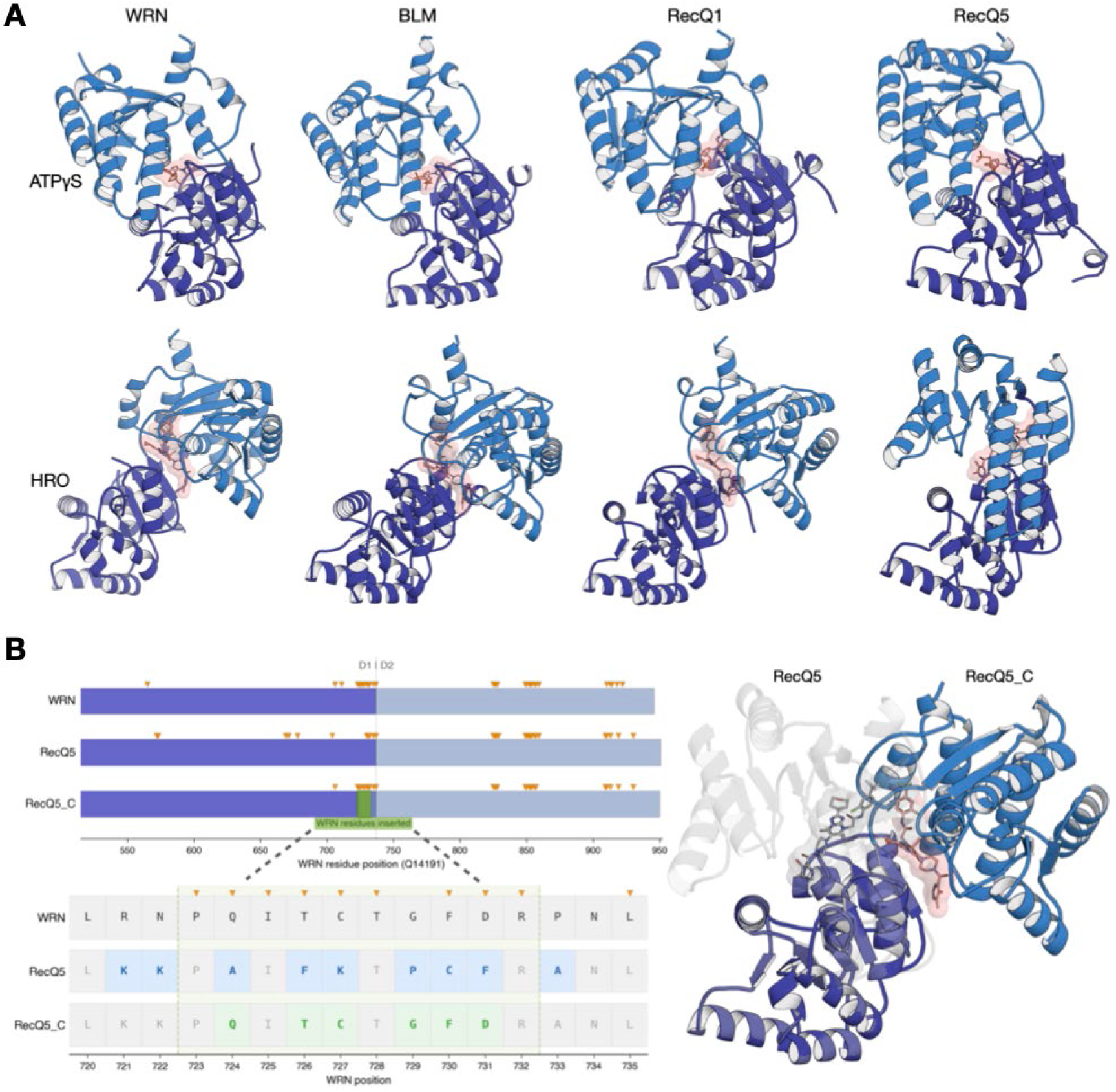
Finetuned model generalizes ligand-dependent conformational switching across RecQ-family helicases in a sequence-dependent manner. A Boltz-FT produces ligand-specific predictions for other RecQ helicases. Werner (WRN), Bloom (BLM), and RecQ1 are each correctly modeled in both their ATPγS-bound active conformation (top) and HRO761-bound-like inactive conformation (bottom), showing the same domain closure and pocket formation observed in WRN. For RecQ5, however, the finetuned model fails to fully adopt the closed state with HRO761 (bottom right, remaining partially open). B Sequence alignment and mutational chimera experiments reveal that the learned allosteric switch is conditional on binding-site sequence. Key WRN/BLM/RecQ1 pocket residues (green) are absent in RecQ5 (orange). Introducing six WRN-derived residues into RecQ5 (creating a RecQ5_C variant) enables Boltz-FT to predict the HRO761-bound closed conformation for RecQ5, whereas reciprocally substituting the corresponding RecQ5 residues into BLM or RecQ1 (BLM_C, RecQ1_C) prevents those helicases from switching to the inactive state. This demonstrates that the finetuned model’s ability to predict the ligand-induced closed conformation is explicitly dependent on the presence of specific sequence features in the binding site.

## Discussion

Our findings demonstrate that protein-ligand co-folding models can be rapidly adapted to capture ligand-induced conformational states that are absent from public structural databases. By finetuning Boltz-1 on ten WRN helicase structures, we enabled the model to predict both a previously unseen allosteric pocket and the large (∼180°) domain rotation induced by HRO761, while preserving its accuracy on the canonical ATP-bound conformation. This reveals that the limitations of co-folding models on dynamic systems stem from training data biases, not fundamental architectural limits, which can be overcome by systematic data generation, as recently suggested^16^. Co-folding networks can be viewed as strong structural priors, updatable as new ligand-bound states emerge from academic labs, drug discovery programs, or community efforts such as OpenBind.

From a practical standpoint, our results illustrate a replicable recipe for injecting new mechanistic knowledge into a foundational co-folding model. The essential components are a small set of carefully chosen structures capturing the desired ligand-stabilized state, at least one example of the alternative conformation (e.g., ATP-bound) in the finetuning mix to preserve dual-state competence, and parameter-efficient adaptation (e.g., low-rank updates in Pairformer layers) to imprint the new behavior without overfitting or catastrophic forgetting. In our case, fewer than 1% of model parameters and ten structures were sufficient to teach a new binding site coupled to a major conformational switch. Notably, even training on chemically dissimilar, weak-potency compounds sufficed to predict the binding mode of a mature drug candidate, indicating that early-stage structural data from a discovery program can recalibrate foundation models without requiring optimized chemical matter.

Beyond WRN, the learned conformational switching behavior transfers across related proteins in a binding site sequence-dependent manner. Although HRO761 itself does not bind to other RecQ helicases with appreciable affinity - reflecting deliberate optimization for target selectivity - such allosteric mechanisms are often conserved within protein families^17^. The mutational experiments demonstrate that the finetuned model learned a sequence-dependent rule coupling binding-site residue identity to conformational outcome. The allosteric transition is triggered only when the requisite residues are present, indicating that finetuning encoded transferable structural logic rather than WRN-specific memorization utilizing ligand information solely as labels for protein conformations. This suggests that analogous ligand-dependent transitions in other target classes (for example, GPCRs switching between active and inactive states, or regulatory enzymes with autoinhibited versus open forms) could similarly be addressed, provided structural exemplars of the alternate state exist.

Our finetuning strategy, however, operates within clear boundaries. The finetuned model selects among discrete end states rather than modeling continuous transitions. This limitation suggests an opportunity for integration with physics-based or hybrid approaches. Molecular dynamics simulations and emerging generative dynamics models excel at sampling intermediate conformations and transition pathways^18^, whereas co-folding networks rapidly identify plausible end states from sequence and ligand cues, possibly interpolated from structurally related family members^19^.

A further nuance is that our finetuning improved structural predictions but did not directly address ligand potency or selectivity. The adapted model learned where and how HRO761 binds in geometric terms, but not how strongly: our training objective was purely structural, incorporating no binding-affinity data. Nonetheless, recovering the correct ligand-stabilized conformation and binding pose is a necessary precondition to build robust 3D-based affinity prediction on top of such models, because erroneous geometry undermines any downstream scoring function or free-energy calculation. In practice, our approach can be paired with physics-based scoring or augmented with activity-labeled structures to translate structural accuracy into quantitative binding energies and rank-ordering of compounds.

Overall, these results indicate that co-folding models’ poor performance on dynamic systems is primarily a symptom of insufficient task-specific adaptation, rather than an inherent inability to capture complex protein–ligand interactions. Furthermore, our findings imply that despite recent findings around the limited performance of co-folding models on unseen target ligand pairs there is a potential path forward by generating representative datasets addressing these gaps and finetune foundational models to enable accurate pose prediction as prerequisite for downstream tasks, such as affinity predictions^20^.

## Methods

### Overview

Boltz-1 is a diffusion-based biomolecular structure prediction model comprising approximately 592 million parameters and follows the architectural paradigm introduced by AlphaFold3.

The model operates on two complementary learned representations: a single (per-token) representation *s* ∈ ℝ^N×384^, where each row *s*i encodes the local biochemical and structural context of token *i*, including residue identity, chemical properties, evolutionary conservation, and, as the network deepens, the token’s inferred spatial environment and interaction propensity; and a pairwise representation *z* ∈ ℝ^N×N×128^, where each element *z*ij encodes the learned relationship between tokens *i* and *j*, including relative positional encoding, co-evolutionary signal from the MSA, and - through iterative refinement - inferred inter-residue distances, orientations, and contact likelihoods. Together, *s* and *z* form a sufficient representation from which the model generates 3D atomic coordinates via diffusion.

The model consists of three key modules: the trunk, the confidence module, and the diffusion-based structure module. The trunk begins with an input embedder that initializes *s* and *z* from atomic features, residue-type encodings, relative-position embeddings, and bond connectivity, followed by an MSA module that integrates co-evolutionary information from multiple sequence alignments into *z* via outer-product mean and pair-weighted averaging operations, and a 48-block Pairformer that iteratively refines both *s* and *z* through triangle operations that enforce geometric consistency in *z* and pair-biased self-attention that transfers pairwise information into *s*. The diffusion-based structure module then generates atomic coordinates through iterative denoising conditioned on the refined *s* and *z*. The confidence module combines trunk-derived representations with dedicated confidence heads to predict per-residue and per-pair quality estimates, including pLDDT, PDE, and PAE, throughout training.

The Pairformer and confidence heads are the primary targets for finetuning because the Pairformer governs what the model learns about inter-residue geometry and structural contacts, whereas the confidence heads determine how accurately the model scores its own predictions for a specific target class.

### Finetuning strategy

We employed Low-Rank Adaptation (LoRA)^2^ to finetune the Pairformer module while keeping the majority of pretrained weights frozen. For a pretrained weight matrix 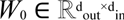 LoRA introduces a low-rank update,

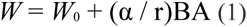

where 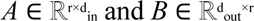 are the trainable low-rank matrices, *r* is the rank (a hyperparameter controlling adaptation capacity), and α is a scaling factor. Matrix *A* is initialized with a Kaiming uniform distribution and *B* with zeros, ensuring that Δ*W* = 0 at initialization and the model reproduces pretrained behavior before any gradient updates. During the forward pass, the adapted linear layer computes,

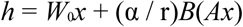

where the frozen pretrained computation W₀x proceeds without gradient tracking and only the low-rank matrices A and B receive gradient updates.

### Choice of rank

The LoRA rank *r* determines the adaptation capacity by setting the number of trainable parameters added to each modified linear projection. For a projection with input and output dimensions *d*_in_ and *d*_out_, LoRA adds two trainable matrices, contributing *r*(*d*in + *d*_out_) parameters. In the square case where *d*_in_ = *d*_out_ = *d*, this becomes 2*rd*, compared with *d*^2^ parameters in the full weight matrix. Thus, relative to full finetuning of the projection, LoRA reduces the parameter count by a factor of *d*/(2*r*).

For the Boltz-1 Pairformer with *d*_s_ = 384 and *d*_z_ = 128, we evaluated ranks *r* ∈ {8, 16, 32}, corresponding to 1.00%, 1.96%, and 3.84% trainable parameters respectively in the LoRA-only configuration.

### Finetuning Targets Pairformer Module

Each of the 48 Pairformer blocks contains two sequential processing stacks that are executed in order - the pairwise stack first, followed by the sequence stack with residual connections throughout.

### Pairwise Stack

The pairwise stack updates the pair representation **z** in ℝ^N×N×128^ through five sub-modules, each applied as a residual:

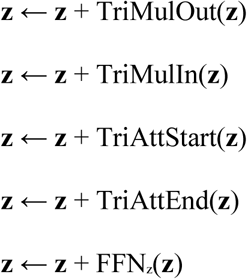

Triangle multiplication (outgoing and incoming) enforces geometric consistency by aggregating over intermediate residues. For the outgoing variant, for each pair (i,j),

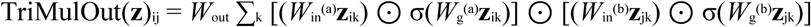

where *W_in_*^(a)^, *W_in_*^(b)^ ∈ ℝ^128×128^ are input projections, *W_g_*^(a)^ and *W_g_*^(b)^ are gating projections, and *W*out ∈ ℝ^128×128^ is the output projection. LoRA is applied to these projections, yielding 20,480 trainable parameters per triangle multiplication sub-module at rank *r* = 16.

Triangle attention (starting-node and ending-node variants) performs self-attention along the rows and columns of *z*, respectively. For the starting-node variant with 4 heads and head width 32:

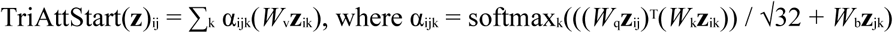

LoRA is applied to all five projections *W*_q_, *W*_k_, *W*_v_, *W*_g_, and *W*_o_ in ℝ^128×128^, plus the bias linear layer *W*_b_ ∈ ℝ^128×4^, yielding 22,592 trainable parameters per triangle attention sub-module.

The transition FFN applies a gated feedforward network to each pair embedding independently,

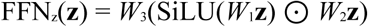

LoRA is applied to all three projections, yielding 30,720 trainable parameters.In total, the pairwise stack contributes 116,864 LoRA-trainable parameters per block across its five sub-modules.

### Sequence Stack

The sequence stack updates the single representation **s** in ℝ^N×384^ through pair-biased self-attention followed by a transition FFN.In the *AttentionPairBias* module (16 heads, *d*_h_ = 24), the attention computation is:

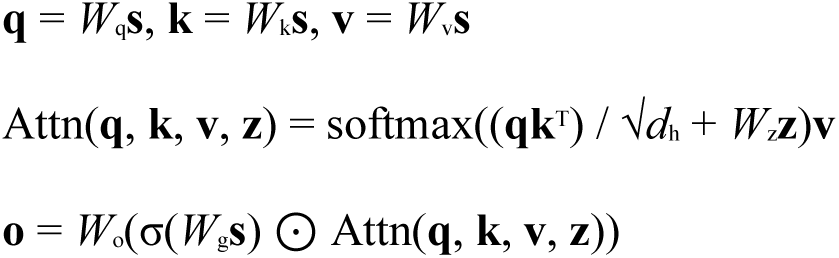

where σ denotes the sigmoid function, ⊙ is element-wise multiplication, and *W*z projects the pairwise representation **z** into per-head attention biases via a *LayerNorm* followed by Linear(128 → 16). LoRA is applied to all five linear projections *W*_q_, *W*_k_, *W*_v_, *W*_g_, and *W*_o_ in ℝ^384×384^, each contributing 2*r* × 384 trainable parameters. The pairwise bias projection *W*_z_ is not adapted because it is an *nn.Sequential* module. With the LoRA-only configuration (no adapter MLP), this yields 5 × 2*r* × 384 trainable parameters per attention block (e.g., 61,440 at *r* = 8).

The sequence transition FFN has the same gated architecture as FFNz, but operates at *d*s = 384 with intermediate dimension 1,536:

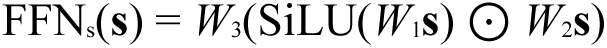

where *W*_1_, *W*_2_ ∈ ℝ^384×1536^ and *W*_3_ ∈ ℝ^1536×384^. LoRA on these three projections adds 92,160 parameters.

The sequence stack contributes 449,664 LoRA-trainable parameters per block (at *r* = 16), dominated by the attention module (357,504) due to the larger embedding dimension *d*s = 384 versus *d*z = 128 in the pairwise stack.

This module is the critical target for finetuning because *AttentionPairBias* is the sole pathway through which learned pairwise geometric constraints (**z**) modulate per-residue features (**s**).

Adapting *W*_q_ and *W*_k_ modifies which residue pairs the model attends to, while adapting *W*_v_ modifies what information is extracted from attended residues. Simultaneously, LoRA on the pairwise stack sub-modules adapts how the model enforces triangle inequality constraints and aggregates geometric information across residue triplets enabling domain-specific refinement of the distance/relationship map that governs structure prediction.

### Confidence Heads

The confidence heads consist of four linear projections (without bias) that map the refined representations to binned quality predictions. These heads (36,352 parameters total) are fully unfrozen during finetuning rather than LoRA-adapted, as they are small terminal projections whose output distributions must recalibrate for the target domain. The confidence loss gradients propagate backward through the frozen confidence module pairformer into the trunk pairformer LoRA adapters, jointly optimising structural representations and quality scoring.

## Author contributions

R.G. performed experiments and data analysis. C.S. and F.S. conceived and supervised the project. All authors contributed to writing the manuscript.

## Competing interests

All authors are employees and/or shareholders of Novartis Pharma.

## Supplementary Information

### Results

#### Overview of externally reported WRN conformations

An independent fragment-based screen (Palte et al., Nat. Commun. 2026) identified multiple ligand-stabilized WRN conformations beyond the canonical ATP-bound state. Using a truncated WRN construct, that study solved ten crystal structures (PDB 9MJS, 9MJT, 9MJU–9MJZ, 9MK0, 9MK1) spanning fragment hits to optimized leads. These structures revealed two new inactive-state categories: Form D, induced by several fragments and early compounds (e.g., PDB 9MJT, 9MJU), which closely resembles the HRO761-bound Form B (sharing nearly all pocket-defining residues despite a subtle D2-domain offset); and Form E, observed with an advanced inhibitor (PDB 9MK0), which engages the same interdomain pocket but rotates the D2 domain ∼90° relative to Form D. These new structures highlight WRN’s conformational plasticity and provided a stringent out-of-distribution test set for our model (see Supplementary Table 3 for a full mapping of WRN forms and PDB IDs).

**Supplementary Fig. 1:**
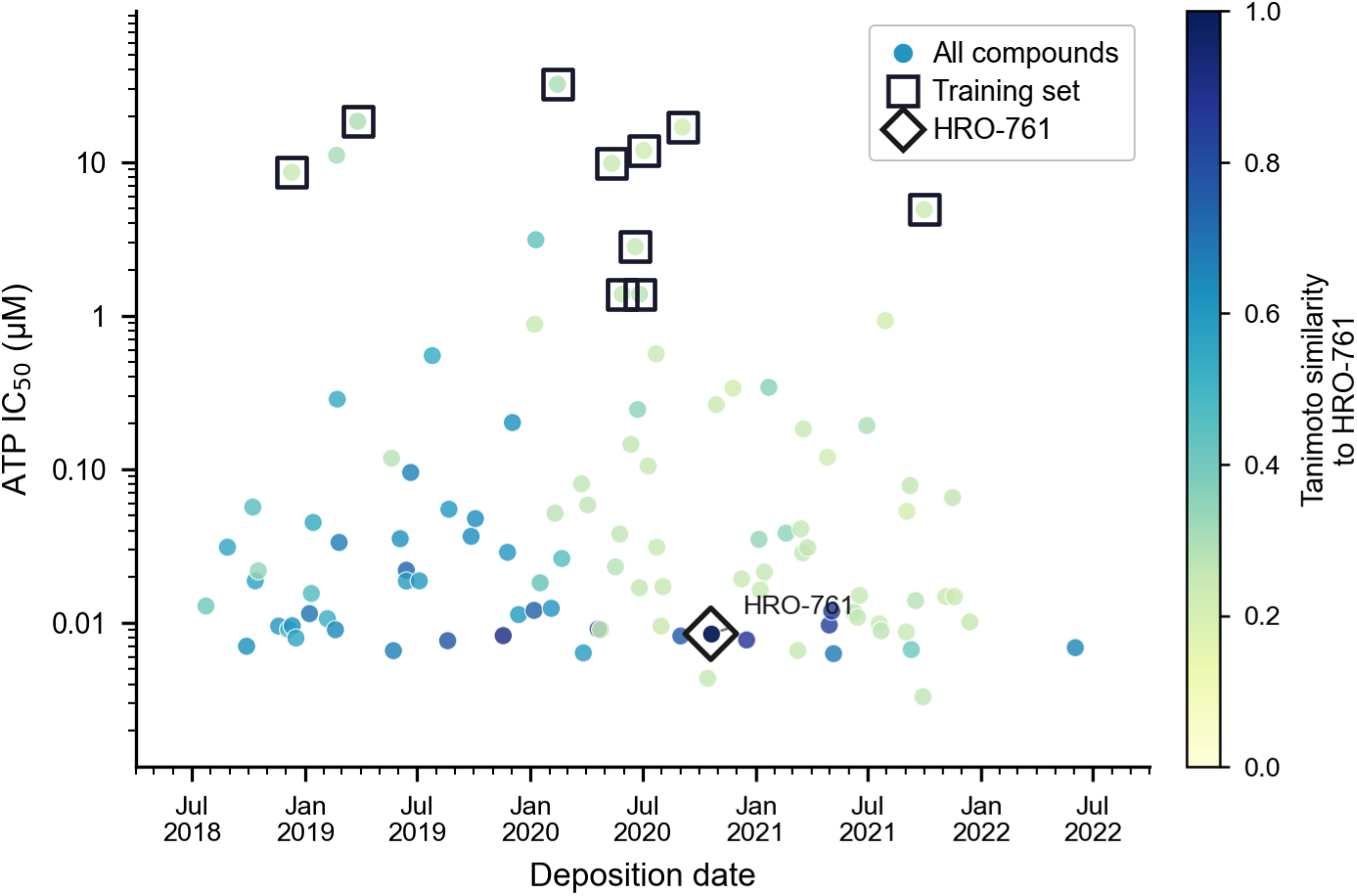
Time distribution and chemical diversity of collected WRN protein–ligand complexes. Scatter plot of ATP IC₅₀ values (µM, log scale) versus deposition date, showing all WRN– ligand complexes gathered over the project timeline. Each point represents one complex; point color encodes Tanimoto similarity to the reference inhibitor HRO761 (0–1 scale, colored from yellow for low similarity to blue for high similarity), illustrating the range of ligand chemotypes. Points outlined with black squares indicate complexes included in the model training set (dissimilarity split), and the HRO761-bound complex is marked by a diamond symbol.

**Supplementary Fig. 2:**
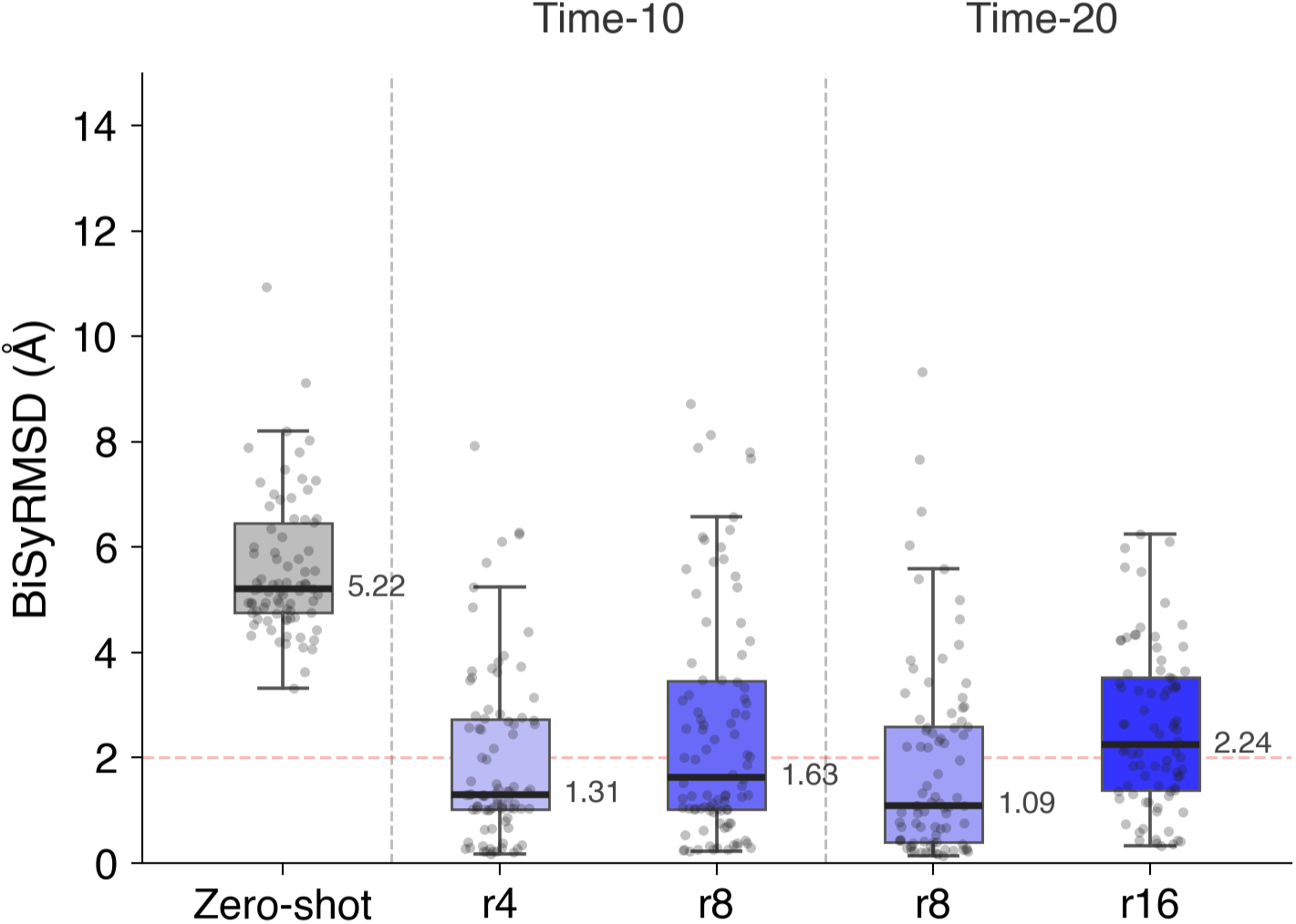
Ablation analysis of LoRA parameters under time-based data splits. Distribution of biSyRMSD values (Å) for HRO761-bound WRN predictions across different training configurations. Results are shown for the zero-shot model and for fine-tuned models with varying LoRA ranks (r), evaluated using a time-based split with either the first 10 (Time-10) or first 20 (Time-20) structures used for training.

**Supplementary Fig. 3:**
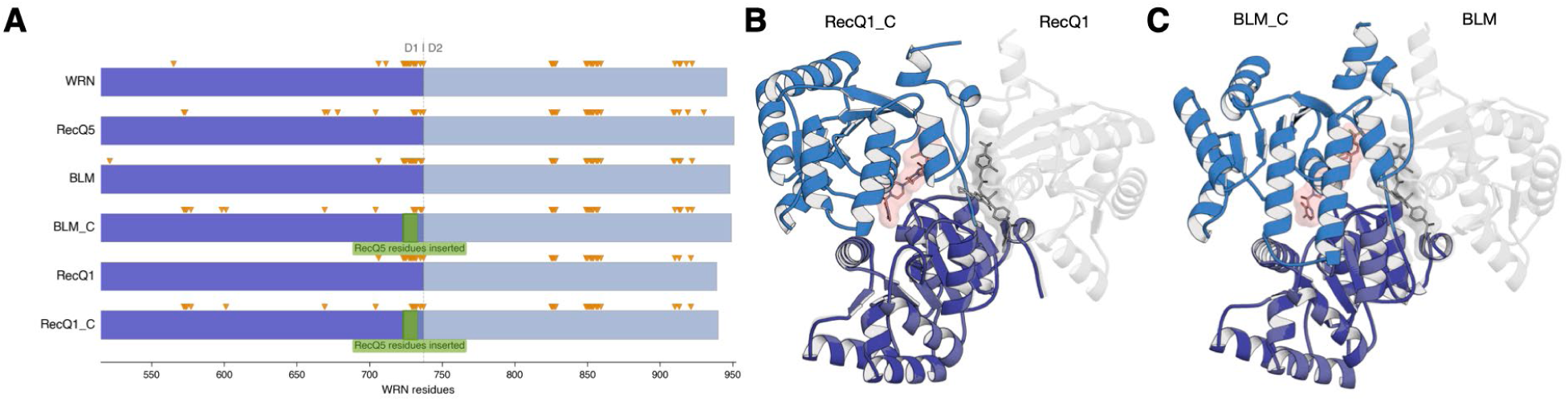
Sequence swaps in RecQ helicases reveal the binding-site sequence dependence of ligand-induced domain closure. **A** Domain schematic of the RecQ-family core helicases (WRN, RecQ5, BLM, RecQ1) and two chimeric variants (BLM_C and RecQ1_C) in which six RecQ5-derived residues have been inserted at the D1–D2 interface (green boxes). The vertical dashed line marks the D1– D2 domain boundary, and orange arrowheads indicate positions where RecQ5 lacks key binding-site residues present in WRN, BLM, and RecQ1. **B** Predicted HRO761-bound structures for wild-type RecQ1 (blue) and its chimeric variant RecQ1_C (RecQ5-sequence insertion, white). The fine-tuned model correctly predicts a fully closed D1–D2 interface for RecQ1 (encapsulating HRO761 in a pocket, pink highlight), whereas RecQ1_C remains partially open and fails to form the closed pocket despite the ligand. **C** Equivalent predictions for wild-type BLM (blue) and BLM_C (white), showing that replacing BLM’s binding-site residues with the RecQ5-derived sequence prevents the inhibitor-induced domain closure. These results demonstrate that the model’s ability to predict the closed, HRO761-bound conformation is abrogated by the RecQ5-like sequence, underscoring the sequence-dependent control of the allosteric conformational switch.

### Methods

#### Training Configuration

Training was performed on a single NVIDIA H100 GPU (80 GB) using PyTorch 2.8 with 32-bit precision.

Key hyperparameters:

The learning rate in Boltz-1 followed the AlphaFold3 schedule with linear warmup to 1.8 times 10^{−3} over 1,000 steps, followed by stepwise decay (factor 0.95 every 50,000 steps).

The diffusion process used rho = 7 for the noise schedule with coordinate augmentation and alignment-based reverse diffusion enabled. Symmetry correction was applied during confidence loss computation to handle symmetric ligand/chain placements.

**Table 1.**
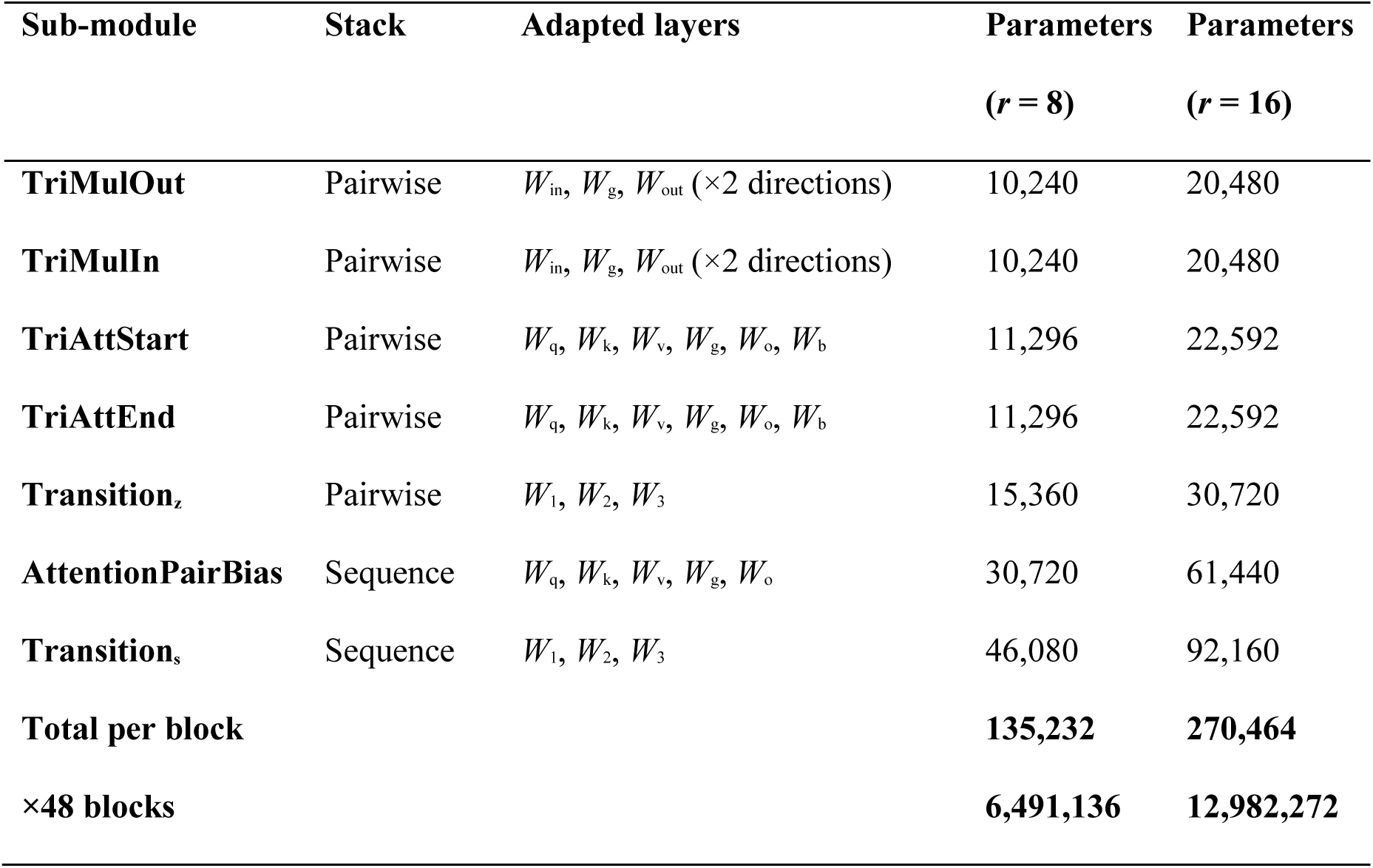
LoRA-adapted sub-modules in each Pairformer block, with stack assignment, adapted layers, and trainable parameter counts for ranks *r* = 8 and *r* = 16.

**Table 2.**
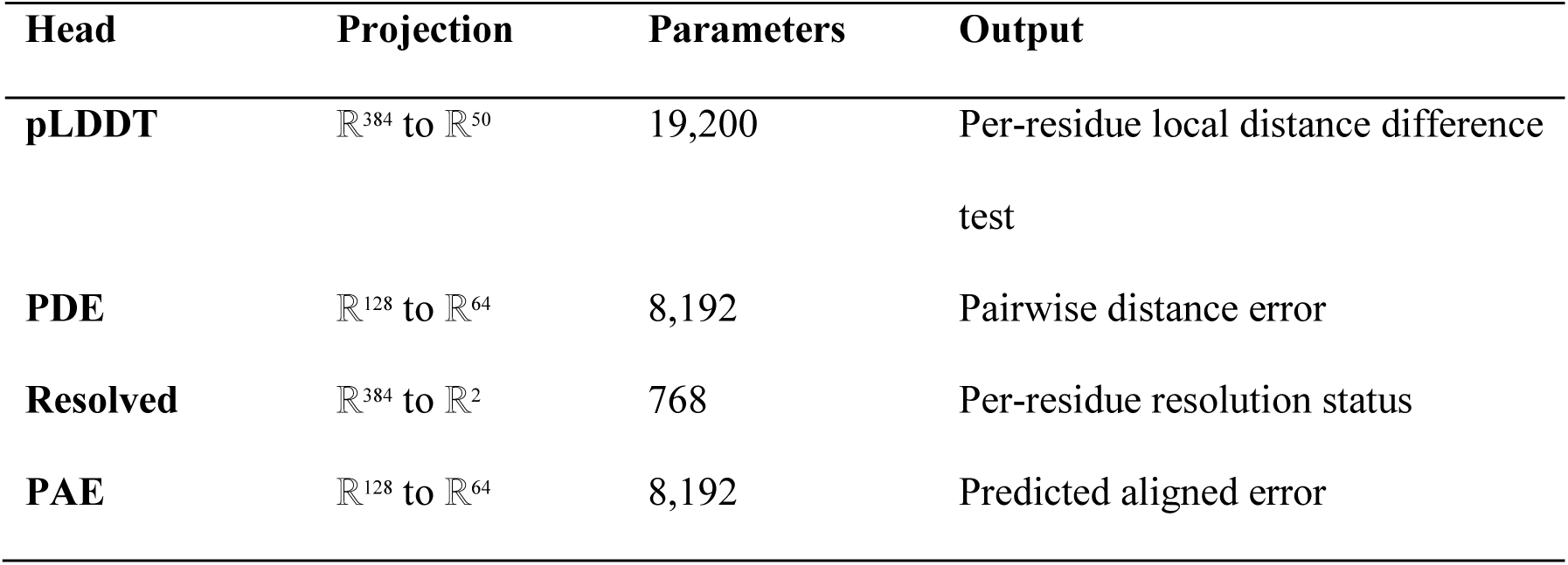
Confidence heads used during finetuning, with projection dimensions, parameter counts and predicted outputs.

| Head | Projection | Parameters | Output |
| --- | --- | --- | --- |
| **pLDDT** | $\mathbb{R}^{384}$ to $\mathbb{R}^{50}$ | 19,200 | Per-residue local distance difference<br>test |
| **PDE** | $\mathbb{R}^{128}$ to $\mathbb{R}^{64}$ | 8,192 | Pairwise distance error |
| **Resolved** | $\mathbb{R}^{384}$ to $\mathbb{R}^2$ | 768 | Per-residue resolution status |
| **PAE** | $\mathbb{R}^{128}$ to $\mathbb{R}^{64}$ | 8,192 | Predicted aligned error |

**Table 3.**
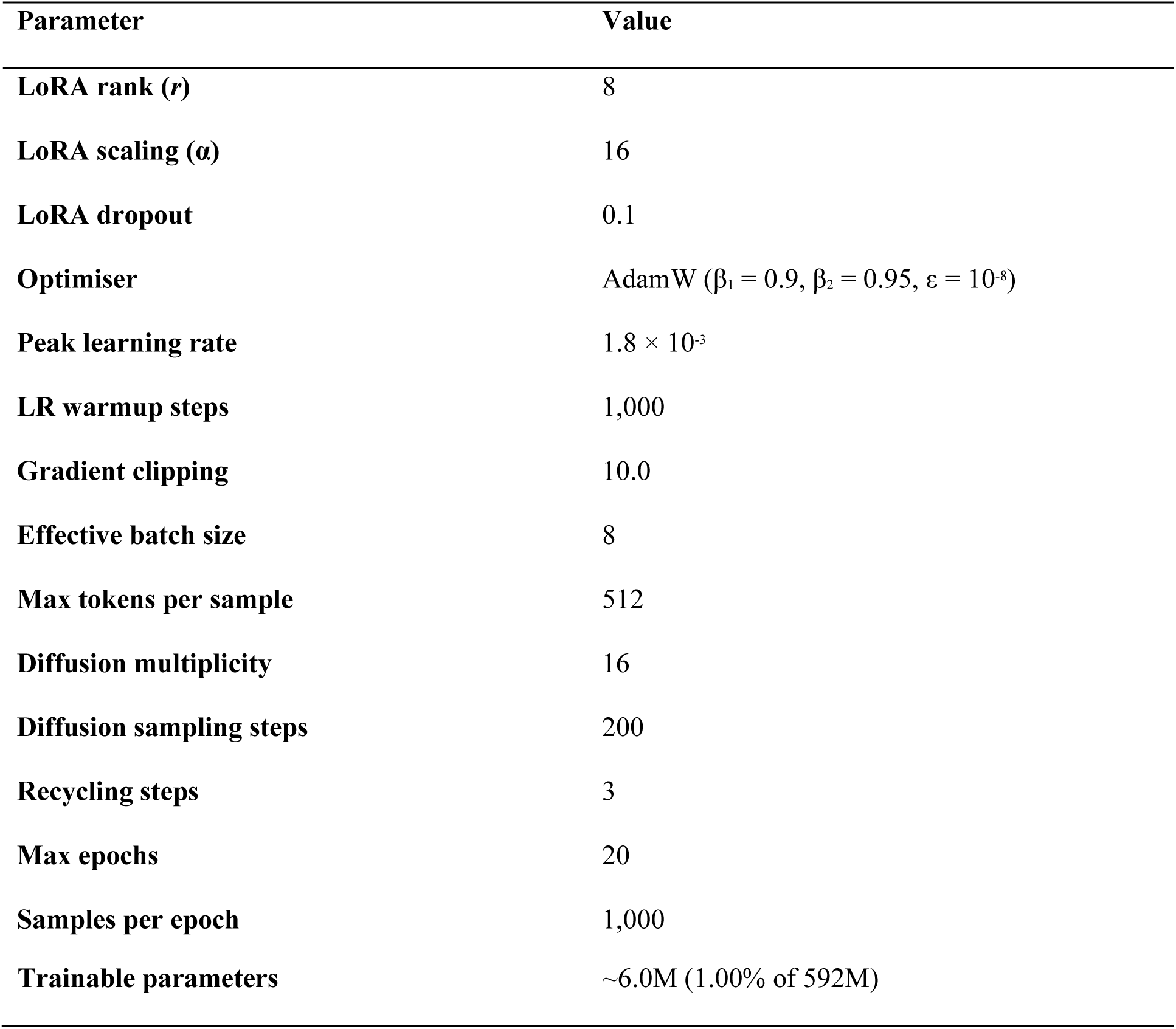
Training configuration used for LoRA finetuning of Boltz-1 on the target-specific structure prediction task.

### Loss Functions

The total training loss is a weighted combination of three terms as proposed in Boltz-1,

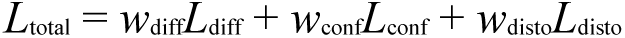

with weights *w*_diff_ = 4.0, *w*_conf_ = 3 × 10^-3^, and *w*_disto_ = 3 × 10^-2^ and we use the default weights for finetuning.

## Notes

### Competing Interest Statement

The authors have declared no competing interest.

